# Contact-dependent regulation of UV-B/C-induced cell fate by neighbouring intact cells

**DOI:** 10.64898/2026.02.11.704850

**Authors:** Jelena Budimir, Nikola Pavlović, Andrea Gelemanović, Vanda Juranić-Lisnić, Swetlana Sperling, Milena Ninković, Miroslav Radman, Katarina Trajkovic

## Abstract

High-energy UV light from the UV-B and UV-C ranges induces severe cellular damage that leads to oxidative stress, cellular senescence, and apoptosis. Most studies of such UV-induced phenotypes have been performed in homogeneous cell cultures where all cells were subjected to comparable levels of damage. However, in physiological conditions, UV exposure generates heterogeneous cell populations in which damaged cells coexist with intact neighbors. How such cellular context influences UV-induced outcomes remains insufficiently understood.

Here, we examined the effects of intact neighbouring cells on mammalian cells exposed to combined UV-B and UV-C radiation. We show that the presence of intact cells enhances apoptotic progression and clearance of cells treated with high doses of UV, while having little effect the cells exposed to low and moderate doses. Transcriptomic profiling revealed that UV-treated cells grown in co-culture with intact neighbours exhibit a markedly attenuated transcriptional response to UV exposure, including reduced activation of oxidative stress and reparatory pathways, compared to UV-treated cells grown in monoculture.

These effects required direct cell–cell contact and were not mediated by diffusible factors, gap junctions, or tunneling nanotubes. Instead, co-culture conditions were associated with extensive changes in ligand–receptor gene expression profiles, indicating altered intercellular communication in response to UV damage.

Our findings demonstrate that UV-induced cellular outcomes are strongly shaped by the surrounding cellular environment and identify a contact-dependent, non-cell-autonomous layer of regulation that influences the resolution of UV-induced damage. These results have implications for understanding tissue-level responses to UV exposure in photodamage, photoaging, and disease contexts.

## INTRODUCTION

UV light damages macromolecules, such as DNA (1) and proteins (2), leading to a range of cellular phenotypes, including oxidative stress (3), premature senescence (4) and cell death (5). The UV spectrum is divided into UV-A (320-400 nm), UV-B (280-320 nm) and UV-C (100-280 nm) regions. The level and type of cellular damage depend on the absorbed dose and wavelength, with shorter wavelengths being more energetically potent and thus more harmful. While UV-B is present in the natural sunlight and poses a risk to skin, UV-C is absorbed by the ozone layer and thus typically not encountered on the Earth surface. However, their combined use can model conditions of severe DNA damage and extreme oxidative stress or mimic exposure in industrial, laboratory or medical settings.

Most of the current knowledge on the cellular effects of UV light was derived from experiments on homogenous cell cultures, where all cells were equally exposed to the stressor. However, UV exposure in tissues generates heterogeneous cellular damage, resulting in the coexistence of severely damaged, mildly damaged, and intact cells. Moreover, a large body of evidence shows that even in tissue culture and much more in physiological conditions, within tissues, same-type cells often display heterogeneous phenotypic and genetic features (6). This might be due to multiple causes, such as uneven exposure to extracellular damaging agents, differences in cell cycle point at the time of stress, or the stochastic nature of somatic mutations that result in somatic mosaicism. Dramatic cell heterogeneity is observed in malignant tissues where genomic instability leads to significant genetic diversification among cells descending from individual cells (7). Of note, cell heterogeneity increases with aging, especially on the transcriptome level (8–10), due to epigenetic alterations. The emergence of senescent cells, each affecting the phenotype of surrounding healthy cells, further contributes to age-related heterogeneity of cellular populations.

In such heterogeneous populations the phenotypes of individual cells can be modulated in a non-cell autonomous manner, depending on the features of the cellular community (11). One example of such non-cell autonomous regulation of cellular phenotypes is the cellular parabiosis, where the presence of healthy cells leads to suppression of aberrant phenotypes in damaged cells (12). In a reverse scenario, cell competition leads to active removal from tissues of the cells with compromised fitness by their fitter neighbors (13). Finally, in the bystander effect, intact cells acquire features of the neighboring irradiated cells (14).

Non-cell autonomous factors mediating these interactions could be transmitted through signaling mediated by intercellular ligand-receptor pairing or via various means of intercellular cargo transfer. Direct transfer of cellular contents between adjacent cells is achieved by means of gap junctions which transfer small molecules - metabolites and ions - or tunneling nanotubes (TNTs) which can transfer proteins and entire organelles. Finally, soluble cargo, such as certain secreted ligands, and extracellular vesicles emanating from cells can be shipped to more remote destinations via the extracellular environment.

In this study, we aimed to characterize the effects of intact cells on the phenotype of cells damaged by combined UV-B and –C light (hereafter UV light) in a heterogeneous cell culture condition. Specifically, we analyzed the impact of intact cells on cell death, apoptotic nuclear phenotype, and cellular transcriptome. Finally, we analyzed the communication tools that may be responsible for the transmission of non-cell autonomous factors. We found that the intact cells promote apoptotic phenotype and reduce transcriptomic response to UV light. This effect of the intact cells was contact-dependent, but it did not rely on GJ and TNTs. However, co-cultures were characterized by dramatically altered ligand-receptor pairing between the two cellular subpopulations.

## RESULTS

In this study, we aimed to investigate whether the phenotype of cells damaged by UV light is affected by the state of the surrounding cells. More specifically, we compared the phenotypes of the damaged cells that were grown in monoculture, i.e., surrounded by roughly equally damaged cells, versus damaged cells co-cultured with intact cells.

### UV light induces dose-dependent apoptosis and senescence in V79 and / or HDF-TERT cells

To establish a baseline for UV-induced cellular responses, we first characterized the dose-dependent effects of combined UV-B and UV-C irradiation on cell survival, senescence, and apoptosis in V79. Application of four doses in the range from 0 to 0.15 J/cm^2^ led to a dose response in the level of damage, as indicated by changes in metabolic activity (Fig. 1A), nuclear fragmentation (Fig. 1B), membrane permeabilization (Fig. 1C), and apoptosis (Fig. 1E, 1F left, Fig. S1). Of note, V79 cells are p53-deficient (15), which suggests that their apoptosis is regulated in a non-canonical manner and that the obtained results (Fig. 1F, left) might be cell line-specific. Thus, we tested the impact of UV light on apoptosis in a different cell line – immortalized human dermal fibroblasts (HDF-TERT) – and obtained similar results (Fig. 1F, right). Unlike other phenotypes, cellular senescence increased at the lower doses (0.015 and 0.03 J/cm^2^) but dropped at the highest dose of 0.15 J/cm^2^ (Fig. 1D), suggesting that the transition to senescence is an active cellular process sensitive to damage by UV light. Together, these results confirm that UV light triggers a spectrum of phenotypes ranging from senescence at lower doses to apoptosis at higher doses, the latter in both V79 and HDF-TERT cells.

**Fig. 1.**
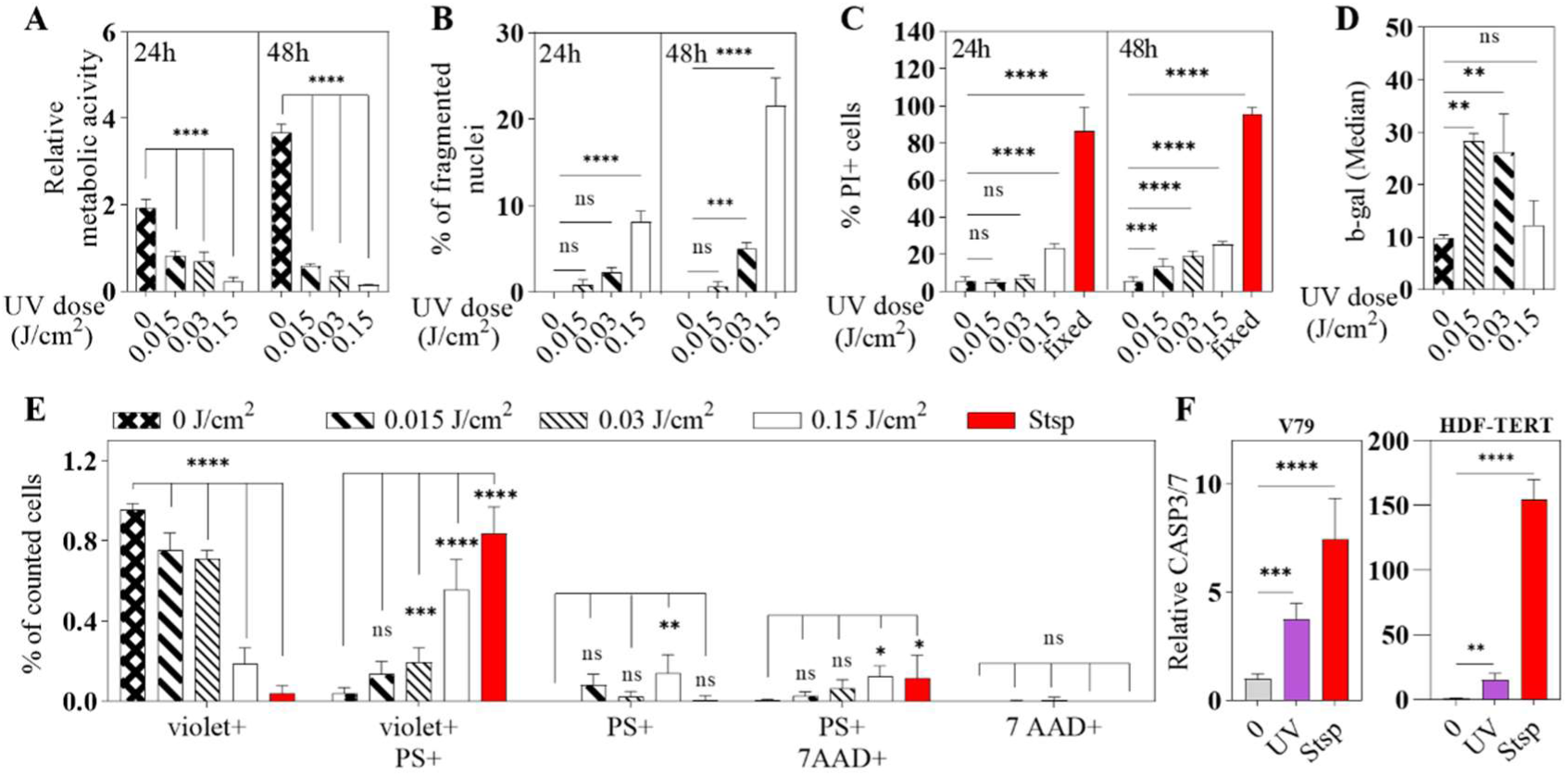
UV light triggers a dose-dependent perturbations in cellular fitness, sencescence and cell death. Cells were treated with a range of doses of UV-light and transferred onto dishes (A, C, D, F), coverslips (B) or chambers for live imaging (E). (A) MTT metabolic activity assay was performed after 4, 24 and 48 h. Relative metabolic activity equals the ratio MTT(24 h) / MTT(4 h). N=16 (technical replicates). (B) Cells were fixed 24 or 48 h after the treatment. Nuclei were stained with DAPI and imaged by fluorescent microscopy. Images of four random microscopic fields were taken and fragmentation of about 200 nuclei per condition was assessed. (C) Cells were stained with PI 24 or 48 h after the treatment. The percentage of PI-positive cells was determined by flow cytometry. (D) Beta-galactosidase activity was determined 24 h after the treatment using flow cytometry. (E) Cells were stained with Cytocalcein Violet (violet+, viable), Apopxin Green (PS+, apoptotic) and 7-AAD (7-AAD+, necrotic) (Apoptosis/Necrosis Kit, Abcam) and imaged live 24 h after the treatment. Images of four or five random microscopic fields were taken, and cells stained with a single or a combination of dyes were counted. Cells treated with 200 μM Staurosporine (Stsp) for 16 h were used as a positive control. Statistical analysis of treated cells in comparison to control (0 J/cm^2^) is shown on the graph. (F) Cells were mock-treated (0) or treated with 0.15 J/cm^2^ UV light (UV). Cells treated with 1 μM Staurosporine for 4 h (Stsp) were used as a positive control. Activation of apoptosis was determined 48 h after the treatment using a CASP3/7 activity assay (Caspase-Glo 3/7, Abcam). Relative CASP3/7 equals the ratio CASP3/7 (UV) / CASP3/7(0).

### Intact cells accelerate removal and apoptotic nuclear phenotypes of UV-damaged neighbours

We next asked whether the state of surrounding cells influences the phenotype of UV-damaged cells by comparing their behavior in monocultures versus co-cultures with intact cells. To answer this question, we plated the equal numbers of fluorescently labelled test cells either in a monoculture or in co-culture with intact V79 cells (Fig. 2A). Since cell density strongly influences the cellular phenotype (16), cell density in the monoculture was adjusted to match the co-culture by growing the fluorescent test cells together with unlabelled, but otherwise equally treated cells (Fig. 2A). The number of the test cells was determined by flow cytometry at 4 h post-plating to assess their initial attachment and at 24 h post-plating to assess the effect of the surrounding cells on retention of the test cells on the plate. The results showed that the presence of intact cells did not affect the relative number of retained test cells treated with 0-0.03 J/cm^2^ (Fig. 2B). However, the relative number of retained test cells treated with the highest dose of UV light was slightly, but significantly lower in co-culture with intact cells compared to the monoculture both at 24 h (Fig. 2B, 2C) and at 48 h (Fig. 2C). An equivalent experiment with HDF-TERT resulted in a similar decrease in the relative number of retained test cells in co-culture with intact cells versus the monocultures at 24 and 48 h post-plating (Fig. 2D). Of note, the initial attachment of the test cells was equal in the co-culture and monoculture (Fig. S2A), indicating that a decrease in the relative number of attached test cells in co-culture is not due to a decrease in the initial attachment. We next asked whether this enhanced removal of the test cells in co-cultures was due to exhaustion of the media in the presence of metabolically more active intact cells. This was not the case, since incubation of the test cells alone in the presence of the media pre-conditioned on confluent intact cells did not affect the number of attached test cells (Fig. S2B). These data show that the presence of the intact cells accelerates the removal of damaged cells from the population. Additionally, the effect of the intact cells seemed to be selective for highly damaged, apoptotic cells.

**Fig. 2.**
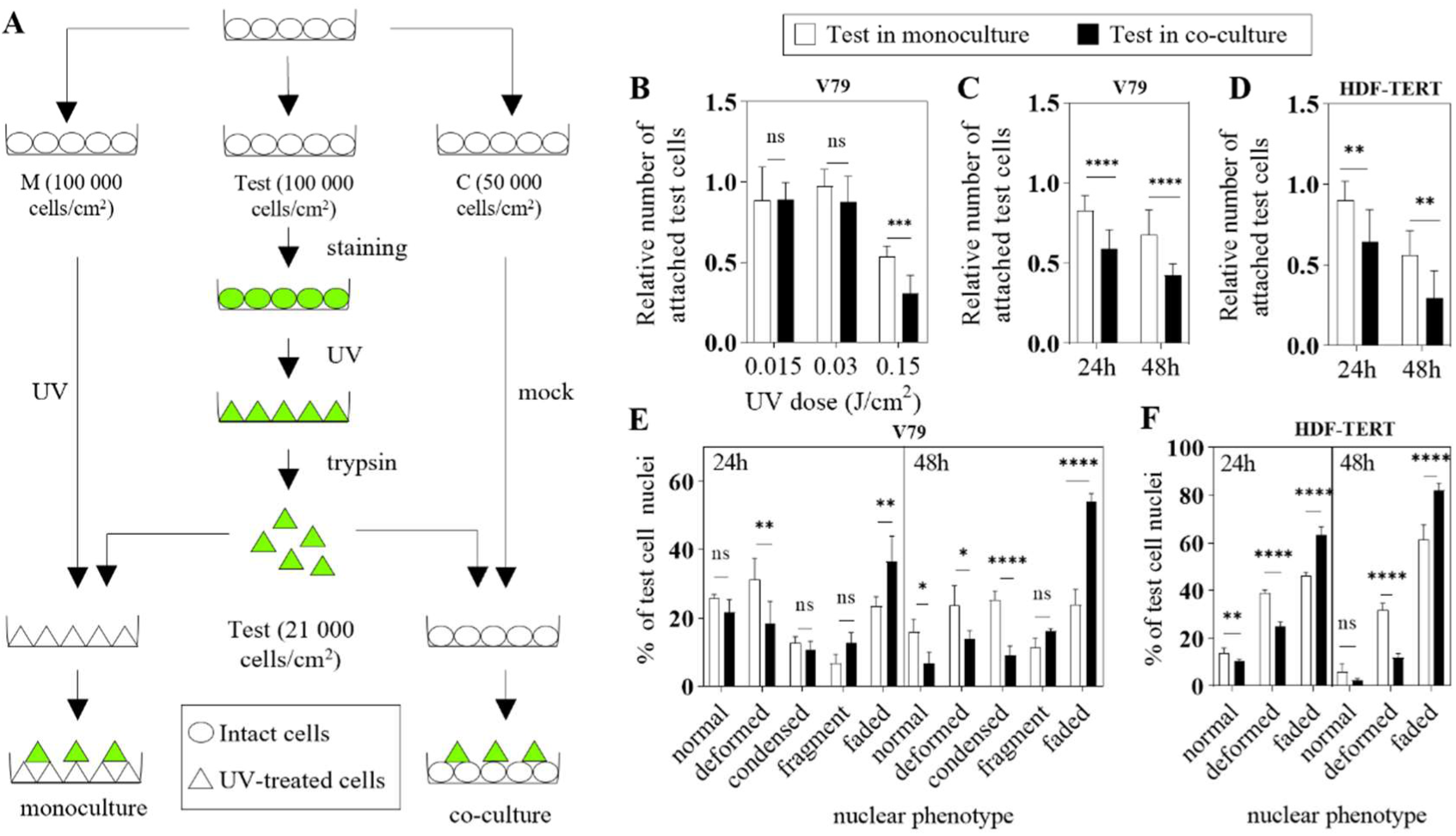
The process of dying in highly damaged cells is influenced by the cellular environment. (A) Fully confluent cells were transferred into three dishes: M (left) for density-matched monoculture, Test (middle) for test cells and C (right) for co-culture. After 24 hours test cells were stained with cytosolic dye (this step was omitted in experiments where test cells stably expressed GFP), treated with UV light, collected and plated into a UV-treated dish ‘’M’’ to form a monoculture and into dish “C” to form a co-culture. The number of attached test cells (N) was determined by flow cytometry 4 h and 24 h after plating. Relative number of attached test cells equals N(time-point) / N(4 h). (B) Number of attached highly damaged V79 cells, but not of the cells with mild damage, is decreased in co-culture with intact cells vs. a monoculture. GFP-expressing test cells were treated with a range of doses of UV light, as explained under (A). (C) The relative number of attached V79 test cells is decreased in co-culture with intact cells compared to a monoculture after 24 h and 48 h. (D) The relative number of attached HDF-TERT test cells is decreased in co-culture with intact cells compared to a monoculture after 24 h and 48 h. (E) Distribution of the nuclear phenotypes of V79 test cells is altered in the co-culture with intact cells compared to the monoculture. Coverslips with monocultures and co-cultures processed as in (C) were fixed after 24 h and 48 h. Nuclei were stained with DAPI and analysed by fluorescent microscopy. About 200 test cells per condition were sorted into one of the five categories based on their nuclear morphology. (F) Distribution of the nuclear phenotypes of HDF-TERT test cells is altered in the co-culture with intact cells compared to the monoculture. HDF-TERT cells were processed and analysed as in (E).

We then tested whether the observed decrease in retention of UV light-treated test cells in monoculture versus co-culture is accompanied by other phenotypic changes. Since these cells undergo apoptosis (Fig. 1E, 1F, Fig. S1), which is characterized by nuclear disintegration, we analyzed the nuclear phenotypes in V79 and HDF-TERT test cells using fluorescence microscopy. Based on nuclear morphology, V79 cells were classified into five groups—normal, deformed, condensed, fragmented, and faded—the latter indicating extensive DNA fragmentation and nuclear degradation typical of late apoptosis (Fig. S3). HDF-TERT cells exhibited similar morphological changes and were categorized into three groups: normal, deformed, and faded. The results showed that the distribution of the cells based on their nuclear morphology was altered in co-cultures versus monocultures, both in V79 and HDF-TERT cells. At 24 h time point the fraction of V79 test cells with faded nuclei was significantly higher in co-culture vs. monoculture, with concomitant decrease in the fraction of cells with deformed nuclei (Fig. 2E). At 48 h time point we observed significantly less test cells with normal, deformed, and condensed nuclei, coupled with a strong increase in the fraction of test cells with faded nuclei in co-culture vs. monoculture (Fig. 2E). Similar changes in distribution of the nuclear phenotypes were observed in HDF-TERT cells – an increase in the fraction of cells with faded nuclei at the expense of other phenotypes both at 24 and 48 h (Fig. 2F). These findings indicate that intact cells selectively exacerbate apoptotic features of highly damaged cells, accelerating their removal from the population.

### Intact cells attenuate the global transcriptional response of UV-treated cells, including oxidative stress pathways

To gain insight into the molecular underpinnings of these phenotypic changes, we performed transcriptome profiling of UV-treated cells in monoculture or co-culture, focusing on stress response and repair pathways. For these experiments, the test cells were incubated in monoculture or co-culture for 12 h to allow initiation of the transcriptomic changes while avoiding massive changes caused by cell death detectable at later time points. This was followed by cell sorting and RNA sequencing of the resulting individual cell populations. Monocultures of intact and UV-treated cells were used as references.

Comparison of the intact and UV-treated reference monocultures showed that the UV treatment caused major transcriptional reprogramming, with 2785 down- and 3612 up-regulated genes (Fig. 3A, Table S1). Gene ontology (GO) enrichment analysis of the differentially expressed genes (DEG) showed upregulation of 111 GO terms and downregulation of 114 GO terms, which were further categorized into biological processes (BP), molecular function (MF), and cellular components (CC) (Table S2). Fig. 3B shows selected GO terms that may be associated with the phenotypes altered in UV-treated cells, as shown in Figs. 1 and 2. These include: (1) upregulation of response to oxidative stress and antioxidant activity, downregulation of mitotic cell phase transition, ribonucleoprotein complex biogenesis and histone H3 demethylase activity (phenotype: decreased metabolic activity, Fig. 1A); (2) upregulation of apoptotic mitochondrial changes and mitochondrial gene expression (phenotype: increased nuclear fragmentation and apoptosis, Fig. 1B, E, F); (3) upregulation of programmed necrotic cell death (phenotype: increased cell permeability, Fig. 1C); (4) downregulation of maintenance of cell number (phenotype: regulation of cell number, Fig. 2), growth receptor binding and Wnt receptor activity (see Fig. 4).

**Fig. 3.**
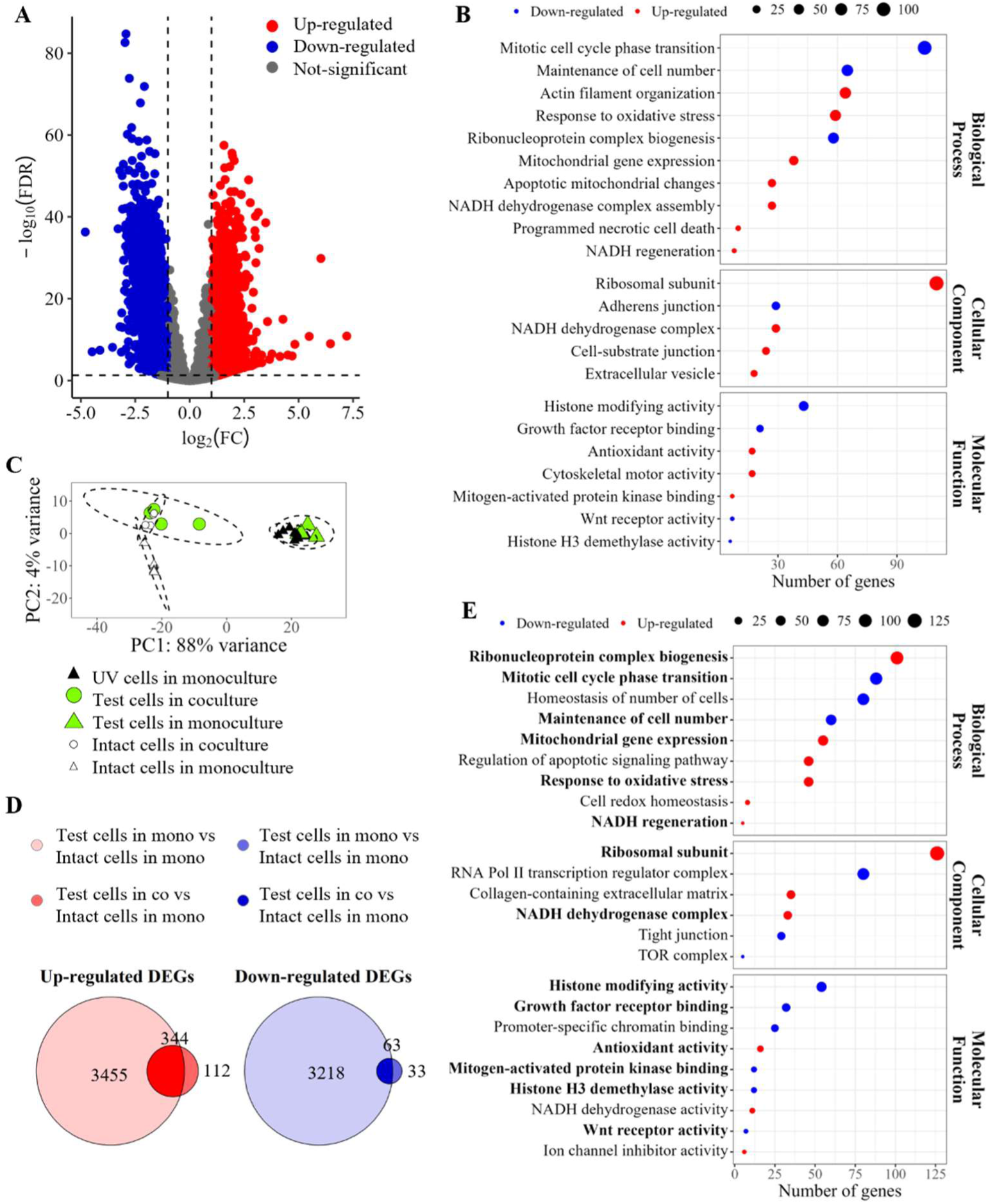
The transcriptional response to UV light is altered in the presence of intact cells. Test cells were incubated in mono- or co-culture for 12 h prior to cell sorting based on green fluorescence. The collected sorted cells were subjected to RNA isolation followed by bulk RNA sequencing. (A) Test cells undergo major transcriptional reprogramming following the exposure to UV light. Differential gene expression (DEG) analysis of intact and UV-treated cells showed that there are 2785 down- and 3612 up-regulated genes after UV exposure (FDR < 0.05, 2-fold change). (B) Selected categories from Gene Ontology (GO) enrichment analysis of DEGs from (A). GO terms in Biological processes, Molecular function and Cellular component (FDR < 0.05, semantic similarity cut-off of 0.5) were selected based on their relevance to phenotypic changes observed after UV exposure. (C) Principal Component Analysis (PCA) of the bulk RNA from sorted samples in quadruplicate showed two distinct clusters: test cells in monoculture (white and green circles, white triangles) and intact cells + test cells in co-culture (green and black triangles). (D) Transcriptional reprogramming of the test cells is different in monoculture vs. co-culture. Venn diagrams of downregulated (blue) and upregulated (red) DEGs from test cells either in monoculture or co-culture and intact cells in monoculture. (E) Selected categories from Gene Ontology (GO) enrichment analysis of DEGs from (D). GO terms in Biological processes, Molecular function and Cellular component (FDR < 0.05, semantic similarity cut-off of 0.5) were selected based on their relevance to phenotypic changes observed after UV exposure. The selected categores that are also present under (B) are shown in bolded letters.

**Fig. 4.**
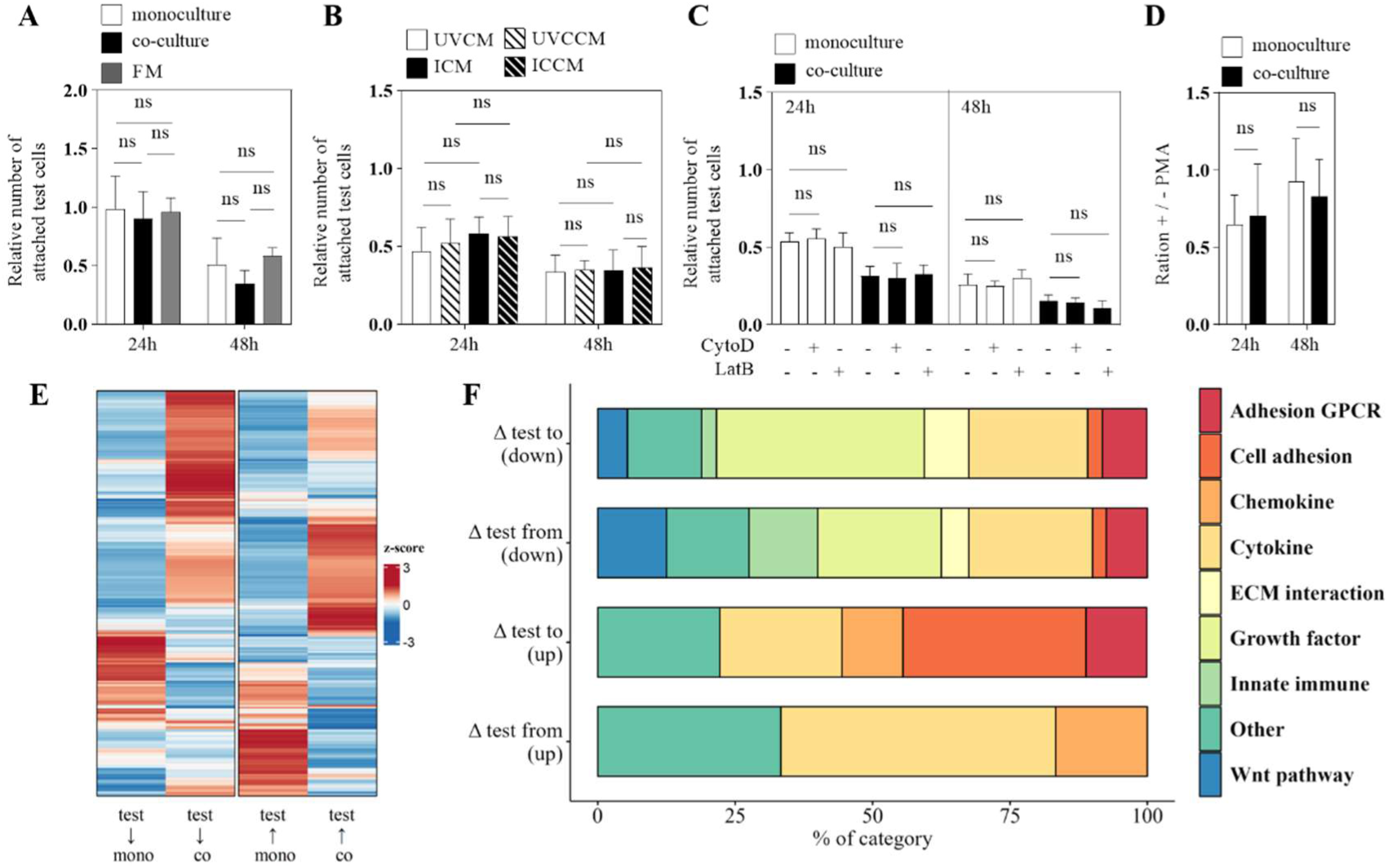
Decrease in number of UV-treated test cells in co-culture vs. monoculture is contact-dependent. (A) The number of test cells remains unchanged in contactless co-culture. Test cells were grown on cell culture inserts with UV-treated cells (monoculture), intact cells (co-culture) or without other cells in fresh medium (FM). (B) Medium conditioned on the challenged intact cells does not lead to a decrease in the number of test cells. Test cells were incubated in the media pre-conditioned for 24 h on a confluent layer of the challenged intact (ICCM) or UV-treated (UVCCM) cells. Media conditioned on unchallenged intact (ICM) or UV-treated (UVCM) cells were used as controls. N=4. (C) Decrease in the number of test cells in co-culture with intact cells does not depend on TNTs. Test cells were plated into density-matched monocultures or co-cultures with the intact cells. The cultures were treated either with the vehicle (DMSO), or with CytoD or LatB. (D) Decrease in the number of test cells in co-culture with intact cells does not depend on GJ. Cells prepared for monocultures and co-cultures were pre-treated for 16 h with vehicle (DMSO) or 50 ng/mL PMA, and further treated equivalently upon addition of the test cells. + / - PMA equals Relative number of attached cells (with PMA) / Relative number of attached cells (without PMA). (E) Scores of ligand-receptor pairs from (↓) and to (↑) test cells in monoculture and co-culture. Analysis was performed using ICELLNET package. (F) Representation of top 10 % ligand-receptor categories with the highest absolute differences between test cells in monoculture and co-culture (Δtest). Enrichment of upregulated (up) and downregulated (down) categories from and to test cells is calculated by subtracting the communication score from and to the test cells in monoculture and co-culture, respectively. Test cells (A-E) were treated with 0.15 J/cm^2^ UV light and their number was determined by flow cytometry after indicated incubation time.

To visualize overall transcriptomic variation among UV-treated test cells grown in monocultures versus co-cultures, principal component analysis (PCA) was performed across all sample conditions (Fig. 3C). Test cells sorted out from the monocultures (green triangle) clustered together with the unlabelled UV-treated cells from the same dish and with the UV-treated reference monoculture (black triangles), as expected. However, test cells from the co-cultures with intact cells (green circle) clustered closer to the intact cells – both to those from the corresponding co-cultures (white circles) and to those from the reference intact monoculture (white triangle).

Comparative analysis of the upregulated and downregulated genes (relative to intact cells) in the test cells in monoculture vs. co-culture showed a dramatically higher number of genes with altered expression in the test cells in monocultures – 3281 vs. 96 down-regulated genes, and 3799 vs. 456 up-regulated genes (Fig. 3D). More than half of the genes with altered expression in co-cultures overlapped with the genes altered in monocultures – 63 for down-regulated and 344 for up-regulated genes (Fig. 3D). Next, Gene Ontology (GO) enrichment analysis was performed on the corresponding differentially expressed genes (DEGs) (Table S3). This analysis showed that about half of the GO categories enriched in UV-treated vs. intact cells (104 out of 225) were enriched in the same direction in test cells in monocultures vs. test cells in co-cultures. GO terms selected based on the relevance for the earlier described phenotypes are shown in Fig. 3E (overlapping categories are highlighted). Of note, enriched GO categories include “response to oxidative stress” and “antioxidant activity”. In conclusion, the transcriptomic analyses reveal that intact cells attenuate the global transcriptional response of UV-treated cells, including oxidative stress–related pathways, aligning their expression profile more closely with that of intact cells.

### Non-cell-autonomous effects of intact cells are contact-dependent and accompanied by altered ligand–receptor signalling networks

Finally, we investigated the mode of communication responsible for the non-cell-autonomous effects, testing the role of factors present in the media (such as diffusible factors or extracellular vesicles), GJ, and TNTs, and mapping changes in ligand–receptor interactions. A search for the cell-cell communication – related GO categories among the DEGs in test cells from mono- vs. co-culture (Table S4) revealed no changes in extracellular vesicle or GJ-associated GO terms, and we were unable to find GO terms in the database that are specifically linked to TNTs. However, 7 GO terms from the Table S4 were linked to receptor-mediated signaling.

We next analyzed experimentally the involvement of various cell-cell communication tools in non-cell autonomous regulation of UV-induced phenotypes. First, we tested whether the influence of the intact cells is mediated by soluble or membrane-bound factors secreted into the media. To that end, we used contactless co-cultures where the test cells were plated on porous inserts placed in a well with a monolayer of irradiated cells, a well with a monolayer of intact cells, or in a well without other cells (Fig. 4A). In the first two cases, the two cell populations were bathing in the same medium, but without physical contact. We observed similar numbers of attached test cells in all conditions after 24 h or 48 h. To address a possibility that a previous direct contact of intact cells with irradiated cells is needed to challenge the intact cells, or that the relevant communication is mediated by vesicles that cannot pass the pores of the inserts, we incubated test cells in the medium pre-conditioned on the intact cells directly confronted with irradiated cells with appropriate controls (Fig. 4B). This approach also showed no significant difference between the conditions at 24 h and 48 h time points. Together, these experiments indicate that direct contact is needed for a reduction of attached test cell number that happened in co-culture with intact cells.

To assess the contribution of TNTs and GJ to the phenotypic differences between the test cells in different cellular environments, the respective communication tools were chemically inhibited. TNTs were dismantled by actin polymerization inhibitors latrunculin A and cytochalasin D, and GJ were inhibited by the phorbol 12-myristate 13-acetate (PMA), which activates protein kinase C (PKC), ultimately leading to phosphorylation of GJ constituent connexin 43. The efficiency of actin inhibitors was confirmed by the disappearance of actin-containing cellular protrusions in the inhibitor-treated samples (Fig. S4A, middle and right image), whereas the efficiency of PMA was evidenced by an increase in the levels of phosphorylated connexin 43 in the PMA-treated sample (Fig. S4B). The test cell number was then compared between the treated and vehicle-treated conditions, both in monocultures and co-cultures. These analyses showed that interference with actin polymerization did not influence the number of test cells in co-cultures or monocultures (Fig. 4C). On the contrary, treatment with PMA reduced the number of test cells both in monocultures and co-cultures at 24 h, but not at 48 h post-plating (Fig. S4C) even though the applied dose of PMA was non-toxic (Fig. S4D). However, this reduction was similar in monocultures and co-cultures (Fig. 4D), suggesting that GJ-mediated communication does not account for the differences in test cell number between the two conditions.

Finally, using the obtained transcriptomic data, we evaluated cell signalling mediated by various ligand-receptor pairs formed between the test cells on one side and the surrounding cells (intact or UV-treated) on the other side. To determine the quantity of such ligand-receptor pairs in mono- and co-cultures, we used transcriptome-based framework ICELLNET v2 (17). ICELLNET v2.2.0 database consists of 1,669 interactions (ligand-receptor pairs), divided into 10 gene families involved in the interaction (cell adhesion, checkpoint, chemokine, cytokine, ECM interaction, growth factor, HLA recognition, innate immune, Notch pathway, and Wnt pathway categories). To estimate the portion of ligand and receptor genes with altered expression in the test cells in co-culture vs. monoculture, we first excluded from the ICELLNET database genes that were not expressed in the test cells, which yielded a total of 363 expressed ligand and receptor genes. Next, these genes were overlapped with the DEGs of test cells in mono- vs. co-culture. This analysis revealed differential expression of 169 ligand and receptor genes (approximately 46.3%) between test cells cultured in mono- versus co-culture conditions. Notably, changes in ligand and receptor gene expression were more pronounced than global transcriptional alterations, as only 28.77% of all genes were differentially expressed. We further assessed ligand-receptor interaction in more detail, taking into account the direction of communication (from or to the test cells) by calculating the communication score between the test cells and the corresponding surrounding cells in mono and co-culture. Heatmap of interaction scores from and to test cells indicates substantial differences in the overall communication profile between test cells and the corresponding surrounding cells in mono- vs. co-culture (Fig. 4E, Table S5). Next, we compared communication scores of each ligand-receptor pair between test cells in mono vs. co-culture and selected the top 10 % pairs with the highest absolute value of the difference in quantity in monoculture vs. co-culture (delta / Δ) (Table S6). Upregulated (in mono- vs. co-culture) ligand-receptor pairs signaling from the test cells included Chemokine, Cytokine, and Other groups, whereas upregulated pairs signaling to the test cells included adhesion GPCR and Cell adhesion in addition to the above three groups. On the other hand, downregulated pairs signaling both from and to test cells belonged to Wnt pathway, Innate immune, Growth factor, ECM interaction, Cytokine, Cell adhesion, Adhesion GPCR, and Other groups (Fig. 4F). The top five most upregulated couples signaling from the test cells to the surrounding damaged cells were GPI/AMFR, LRPAP1/SORT1, TNFSF13B/TNFRSF13C, CXCL16/CXCR6 and TXLNA/STX4, and those signaling in the opposite direction were FLRT1/ADGRL1, ANGPT2/ITGA5+ITGB1, OMG/PTPRS, CCL5/SDC1 and CCL27/CCR10 (Table S6). Together, these experiments and analyses show that the non-cell-autonomous effects of intact cells are contact-dependent and coincide with widespread rewiring of ligand–receptor signaling networks, rather than being mediated by diffusible factors, gap junctions, or tunneling nanotubes.

## DISCUSSION

In this study, we demonstrate that intact cells shape the outcome of UV-induced damage in neighboring cells in a UV dose- and intercellular contact-dependent manner. At high UV doses, co-cultured intact cells accelerated apoptotic clearance and nuclear disintegration of damaged cells, while dampening their transcriptional response to UV, including oxidative stress defenses. These effects were not mediated by diffusible factors, gap junctions, or tunneling nanotubes, but coincided with large-scale remodeling of ligand–receptor signaling networks. Taken together, these findings establish that UV-triggered cellular phenotypes are not solely cell-autonomous but are strongly regulated by direct interactions with intact neighbors, revealing an additional layer of control over damage resolution within cell populations.

Test cells treated by 0.15 J/cm^2^ of UV light were dying by apoptosis (Fig. 1E, F) with a time-dependent reduction in the number of attached cells (Fig. 2B-D). We showed that this reduction in cell number was more prominent in the presence of intact cells at 24 h and 48 h of co-culturing (Fig. 2B-D). This could be due to increased cell detachment, promotion of apoptosis across the irradiated population, or phagocytosis of the apoptotic cells by the surrounding intact cells. Our result is in agreement with studies showing that the professional phagocytes or other cells capable of engulfment, such as fibroblasts, can not only clear the remnants of apoptotic cells, but also contribute to apoptosis by their engulfment machinery (18,19). Furthermore, the surrounding normal cells can actively extrude the apoptotic cells (20).

The observed reduction in the number of attached treated cells co-cultured with intact cells took place only when they were irradiated with a high dose of UV light (0.15 J/cm^2^), which triggered apoptosis (Fig. 2B; 1D-F). On the contrary, intact cells had no effect on the number of attached cells treated with lower doses of UV light (Fig. 2B), which displayed a low level of apoptosis and a high level of senescence (Fig. 1D-F). One possible explanation for this selective effect of the intact cells on apoptotic cells can be found in the “supercellularity” concept described by Rustom (21). He suggested that mildly and moderately stressed cells receive mitochondria via TNTs from their healthy neighbors, which ameliorates the outcome of their stress. However, if the cells are heavily damaged, the intercellular connections are severed, resulting in isolation and removal of the damaged cells from the collective. This concept is consistent with our result that the effect of intact cells on apoptotic cells did not depend on TNTs (Fig. 4B).

Previous research on the effect of intact cells on the phenotype of cells irradiated by UV light is limited and somewhat conflicting. While the work of Wang and Gerdes demonstrated the rescue from apoptosis in co-cultures of UV-treated and intact cells (22), Krzywon and colleagues showed that the surrounding intact cells could both rescue and intensify damage in UV-treated cells, depending on the intact cell type (23,24). Of note, these studies used different cell types, as well as kinds and doses of UV light. Together with our data, the above results suggest that there might be no universal principle in the effect of intact cells on the ones damaged by UV light, which warrants specific studies for various species, cell types, or types of cellular damage.

In this study, we show for the first time that the transcriptome of the cells damaged by the UV light and grown in co-culture with intact cells is more similar to that of the intact cells then of the treated cells in monoculture (Fig. 3C), suggesting that the presence of intact cells attenuated their transcriptomic response to UV light. In particular, their reparatory mechanisms seemed to be poorly active. These include genes involved in response to oxidative stress and antioxidant activity, as well as genes encoding for ribosomal subunits and ribonuclear complex biogenesis which are needed to support the cellular recovery (Fig. 3E). It might be that it is safer and more energy-efficient for a cell population to dismiss a severely damaged cell by inhibiting its inducible protective and rescuing mechanisms, than to invest in its repair and allow for its potentially toxic persistence, for instance undergoing senescence. Moreover, stressed cells express stress ligands due to which the immune cells can attack the tissue; such activation of the immune system may lead to injury of healthy bystander cells (25). Such removal of the damaged cells would make sense, especially under the conditions with a large excess of intact cells, which can proliferate and replace the damaged cells (26).

We show here that, above a tipping point damage threshold, intact cells lead to overall exacerbation of the damage caused by UV light (Fig. 2). This effect of the intact cells resembles cell competition, but maybe more a kind of active tissue cleansing. Another similar example is the case of “assisted suicide” of the stressed germline cells in *C. elegans*, where CEP-1-induced apoptosis of germline cells is positively regulated by the cells in the somatic gonad (26) or in the intestine (27). Of note, in the latter case, apoptosis in one cell type is regulated by a different cell type, whereas in cell competition or the case of tissue sanitation described in our study it is regulated by the same-type cells with different fitness levels.

Our data indicate that the exacerbated phenotype observed in co-culture depends on direct contact between the two cellular subpopulations (Fig. 4A, B). However, we cannot rule out the involvement of diffusible short-lived paracrine factors or ligands and microvesicles that may require high local concentrations to exert their effects. The observed non cell-autonomous regulation in our study did not depend on direct communication tools such as GJ and TNT, as their chemical inhibition did not abolish the effect of intact cells (Fig. 4C, D). These results align with our transcriptomic data analysis (Table S4), which showed no changes in GO terms for microvesicles or gap junctions, and no TNT-specific GO terms were available in the database. Interestingly, other studies have shown that soluble factors participated in non-cell autonomous correction of the phenotype, as in the case where tumor cell proliferation and motility were inhibited by fibroblasts (28). Moreover, exosomes could rescue proteostasis defects (29), and TNT-mediated transfer of mitochondria participated in the rescue from apoptosis (22,30) or of cellular dysfunction (31). On the other hand, GJ, which are essential in metabolic cooperation of the cells, are known both for their ability to propagate and to prevent cell death across the cellular society (32).

Non-cell autonomous regulation of phenotypes can also be mediated by ligand-receptor coupling between adjacent cells. This is the case in some forms of cell competition where cells sense lower fitness of their neighbors via Slit-Robo, Spaetzle-Toll, or Sas-PTP10D interactions (33). Our transcriptomics data show a dramatic difference in the expression of a vast array of ligand-receptor couples in co-cultures of damaged and intact cells vs. monocultures of damaged cells (Fig. 4E). The majority of these couples belong to cytokine, chemokine, growth factor, cell adhesion, innate immune, aGPCR, Wnt pathway and ECM interaction groups (Fig. 4F). Interestingly, a recent article showed that cell-cell signaling can be altered non-cell autonomously in the follicle cells due to reactive oxygen species (ROS) released by the adjacent germ cells (34). This result is consistent with our work since UV radiation also leads to the production of ROS. Here, we provide a list of the most strongly altered ligand–receptor candidates, whose potential roles in fitness sensing or in regulating the phenotype of damaged cells remain to be elucidated, as these questions were beyond the scope of the present study.

In summary, our results show that UV-induced changes in damaged cells are not determined only in a cell-autonomous manner, but are influenced by direct contact with intact neighboring cells. Intact cells promoted the apoptotic progression and clearance of heavily damaged cells while reducing their transcriptional response to UV light, including oxidative stress response and other repair mechanisms. This regulation was contact-dependent and coincided with broad changes in the expression of ligand–receptor pairs. These findings support a model in which healthy cellular environment favors elimination of highly damaged cells over their attempted recovery. Such population-level regulation may represent a tissue-protective strategy relevant to photodamage, photoaging, regeneration after injury or disease context and could point to novel strategies to enhance tissue resilience (44).

## MATERIALS AND METHODS

### Reagents

**General chemicals**: Phorbol 12-myristate 13-acetate (PMA), bromophenol blue, paraformaldehyde, Latrunculin B, protamine sulphate (P4020) and gelatine were from Sigma-Aldrich, USA. Super Signal Pico Substrate, Lipofectamine 3000 and Hoechst 33342 were purchased from ThermoFisher, USA. Tris, NaCl, 3-(4,5-dimethylthiazol-2-yl)-2,5-diphenyltetrazolium bromide (MTT), dimethyl sulfoxide (DMSO), bovine serum albumin (BSA) and milk powder were from ROTH, Germany. Glycerol, HCl and formaldehyde were purchased from Kemika, Croatia. Sodium dodecyl sulphate (SDS) was purchased from Serva, Germany. Tween 20 and β-mercaptoethanol were purchased from Fisher, USA. Staurosporine was from Abcam, UK. RLT buffer was from Qiagen.

**Cell culture reagents**: Dulbecco’s Modified Essential Medium (DMEM) High Glucose with L-Glutamine, phosphate buffer saline (PBS), HEPES, Hank’s Balanced Salt Solution (HBSS), trypsin and foetal bovine serum (FBS) were purchased from Capricorn Scientific, Germany. Cell Proliferation Staining Reagent, Anti-Fade Fluorescence Mounting Medium, 4’,6-diamidino-2-phenylindole (DAPI), Senescence Assay Kit and Apoptosis/Necrosis Assay Kit were from Abcam, UK. CellMask ® Green Actin Tracking Stain and propidium iodide (PI) were from Invitrogen, USA. Caspase-Glo ® 3/7 Assay System was from Promega. All cell culture reagents used for lentivirus production (see below) were purchased from GIBCO, USA. **Antibodies**: rabbit anti-phospho-Connexin 43 (Ser368) Antibody #3511 (used at 1:1000) and goat anti-rabbit IgG HRP (horseradish peroxidase)-linked Antibody #7074 (1:2000) were from Cell Signaling Technology, mouse anti-alpha-tubulin (DM1A) Antibody #62204 (used at 1:1000) and goat anti-mouse IgG (H+L) (DyLight^TM^ 550) Antibody #84540 (used at 1:2000) were from Invitrogen.

### Cell culture

V79 cells (Chinese hamster pulmonary fibroblasts) were obtained from the American Type Culture Collection (ATCC), HDF-TERT cells (human dermal fibroblasts immortalized by overexpression of telomerase) from EVERCYTE GmbH, and HEK 293FT from Thermo Fisher Scientific. All lines were maintained in DMEM with 10 % FBS at 37 °C and 5 % CO2. V79 cells were passaged every 2-3 days at 1:40 dilution, HDF-TERT every 7 days at 1:10 dilution, and HEK 293FT every 3-4 days at 1:10 dilution.

### Generation of stable GFP-expressing V79 cell line

#### Lentivirus production

For the standard transient transfection, HEK 293FT cells were seeded at 4 x 10^6^ cells/ml in a 75 cm² flask containing the following medium: DMEM, 10 % FCS, 1 x sodium pyruvate, 1 x NEAA, 1 x L-glutamine, penicillin/streptomycin and 1 x geneticin. Immediately, lentivirus packaging medium (Opti-MEM reduced serum medium containing 5 % FCS, 1 x sodium pyruvate and 1 x GlutaMAX without antibiotics) was added to the cell mixture at a ratio of 2:1 (v/v). After 2 h, the cells were transfected with the following plasmids (100 µg DNA/75 cm² flask): psPAX2 (#12260, Addgene), pCMV-VSV-G (#8454, Addgene) and p-EGFP (#165830, Addgene) along with Lipofectamine 3000 transfection reagent in a 2:1 (w/w) ratio. The viral supernatant was harvested 72 h after transfection, filtered using a 0.45 µm filter (#MPSCHVU01RE, Merck), concentrated to 1 ml using a Pierce Protein Concentrator with a 10K MWCO (#88517 for 2–6 ml or #88528 for 5–20 ml) and used for transduction immediately.

#### Lentivirus transduction into the target V79 cells

V79 cells were seeded in DMEM with high glucose (4500 mg/l) and 10 % FCS in a 6-well plate at 5 x 10^4^ cells/well. Fresh, concentrated viral supernatant (250 µl/well) and culture medium were added 24 h after seeding. The plates with the lentivirus-infected cells were then centrifuged for two hours at 330 x g (Rotanta 460 RF, Hettich, Germany) at room temperature. After this, protamine sulphate (4 mg/ml in ddH₂O) was added to the medium (1:1000). The cells were expanded with DMEM with 10 % (v/v) FCS. After at least three passages, they were harvested and sorted for GFP fluorescence by flow cytometry (BD FACS ARIA II, San Jose, CA, USA).

### Treatment with the UV light

Fully confluent monolayer of cells was washed two times in PBS and prepared for treatment by adding 0.1 ml of PBS per cm^2^. Irradiation of the cells was performed in the UV irradiation chamber (BS-02, Opsytech Dr Gröbel, Germany) using a combination of UV-B and UV-C light at 2:1 ratio at a dose rate of 5 and 3.5 mJ/s respectively. Immediately after the treatment cells were washed three times in PBS, trypsinized, and processed further as indicated.

### Senescence assay and flow cytometry

Senescence Assay Kit was used to detect senescence-associated beta galactosidase following manufacturer’s instructions. Briefly, trypsinized cells were washed once in PBS and fixed in 4 % formaldehyde for 10 minutes at room temperature (RT), followed by sequential washes in PBS and 10 % BSA in PBS. One half of each sample was incubated for 90 min in the staining probe diluted at 1:500 in the assay buffer. The other half of the sample was used as blank, and it was exposed to the equivalent treatment but without the staining probe. Cells were then washed in 10 % BSA in PBS and resuspended in the same buffer for flow cytometry at a density of approximately 10^6^ cells per mL. The final values were calculated by subtracting blank from the median of each sample.

In co-culture experiments, adherent cells were prepared for flow cytometry from a cell culture plate or insert (Falcon® Permeable Support, 1 µm PET Membrane, Corning) by one wash with PBS, trypsinization and resuspension in 10 mM HEPES/HBSS with 10 % FBS at a density of approximately 600 000 cells per mL.

For propidium iodide (PI) staining, cells collected for flow cytometry as above were washed twice with PBS and stained in 500 µL of 1 µM PI in 20 mM HEPES/HBSS for 10 minutes at a density of approximately 800 000 cells per mL. Cells fixed with 4 % formaldehyde served as a positive control.

For flow cytometry, 200 µL aliquots of the cell suspension were loaded into a well of the 96-well plate and analysed using a Guava easyCyte flow cytometer (Luminex, USA) controlled by guavaSoft software (Luminex, USA).

### Preparation of the conditioned media

Media were conditioned for 24 h on a confluent layer of intact or irradiated cells, clarified by centrifugation at 3 000 rpm for 10 min (Heraeus Megafuge 40R, Thermo Scientific, Massachusetts, USA) and used immediately in experiments. Confronted conditioned media were prepared on a confluent layer of intact or irradiated cells with addition of 21 000 irradiated cells/cm^2^.

### Fluorescent labeling of the cells and microscopy

Labelling of the living cells with Cell Proliferation Staining Reagent was performed following the manufacturer’s recommendation. Briefly, confluent cells were washed with PBS, incubated for 30 minutes at RT in the staining reagent diluted at 1:500 in 20 mM HEPES/HBSS, and washed again three times with PBS.

For the live staining of actin and the nuclei, cells plated on imaging slides (µ-Slide 8 well, ibidi, Gräfelfing, Germany) were labeled for 30 minutes with CellMask ® Green Actin Tracking Stain (1:1000) and Hoechst 33342 (20 µM) nuclear stain in cell culture media supplemented with 10 % FBS. After labeling, cells were washed twice in PBS and imaged in 20 mM HEPES/HBSS.

Apoptosis was detected using Apoptosis/Necrosis Assay Kit following manufacturers’ instructions. Briefly, intact or irradiated cells were washed with assay buffer, incubated for 30 minutes at RT in the assay buffer containing Apopxin Green (1:100), 7-AAD (1:200) and Cytocalcein Violet (1:200), washed twice in the assay buffer and imaged in 20 mM HEPES/HBSS.

Nuclei of the fixed cells were stained with 5 μM DAPI for 10 minutes and washed three times with PBS. For microscopy of the fixed cells the coverslips were rinsed with ddH2O, air-dried and mounted onto the glass slides with a drop of the mounting medium.

Imaging was performed at the fluorescent microscope Axio Observer Colibri 7 (Zeiss, Germany) controlled by Zen BLUE software using UV and blue excitation lines. Images were processed in ImageJ.

### Western blotting

Cells were washed with PBS and lysed directly in 2 x Laemmli buffer (10 % glycerol, 62.5 mM tris/HCl, 2 % SDS, 0.01 % bromophenol blue and 5 % β-mercaptoethanol). 25 µL of the protein lysate was loaded onto a 15 % polyacrylamide gel for SDS-PAGE, followed by transfer from the gel onto a polyvinylidene fluoride (PVDF) membrane (Merck-Millipore) using a wet transfer system (Biorad, USA). The membranes were blocked by shaking for 1 h in 5 % milk diluted in TBS-Tween (20 mM Tris, 150 mM NaCl, 0.05 % Tween 20). The membrane was then incubated overnight at 4 °C with a primary antibody diluted in 5 % BSA in TBS-Tween, washed three times in TBS-T, and incubated for 2 h in the secondary antibody diluted in TBS-Tween. The proteins of interest were revealed either by chemiluminescence or fluorescence. In the case of chemiluminescence, a Super Signal Pico Substrate was used and X-ray films FUJI FPM 100A (Fujifilm, Japan) were exposed to the membranes and developed in Fuji medical film processor (FPM-100A, Fuji Photo Film Co., LTD, Tokyo, Japan). In the case of fluorescence Typhoon™ FLA 9500 biomolecular imager was used (GE Healthcare Life Science) and the images were analyzed using ImageJ (Fiji, Japan).

### Metabolic activity and apoptosis assays

To determine metabolic activity, cells were seeded at the confluency of 100 000 cells/cm^2^. 3-(4,5-dimethylthiazol-2-yl)-2,5-diphenyltetrazolium bromide (MTT) assay was used according to the manufacturer’s instructions. Briefly, the cells were incubated with the MTT-containing medium for 4 hours at 37 °C, and 5 % CO_2_, followed by solubilization of the generated formazan in DMSO. The absorbance was measured at 595 nm using a multi-plate reader (TECAN Sunrise, California, USA).

Activity of apoptosis markers Caspase 3 and 7 was detected using Caspase-Glo ® 3/7 Assay System according to the manufacturer’s instructions. Briefly, 17 500 intact V79 cells, 35 000 irradiated V79 cells or 15 000 HDF-TERT cells per well were seeded in a 96-well plate. After 24 hours, Caspase-Glo ® 3/7 substrate diluted in Caspase-Glo ® 3/7 buffer was added to the cells and incubated for 2 hours at room temperature. Luminescence was detected on Typhoon™ FLA 9500 biomolecular imager (GE Healthcare Life Science) and the images were analyzed using ImageJ (Fiji, Japan).

### Cell sorting, RNA isolation and quality control

Cells (co-cultures and individual cells) were detached as described previously and then passed through 100 µm cell strainer to remove clumps. The cells were then sorted on BD FACSAriaIIu cell sorter (Center for Proteomics, University of Rijeka Faculty of Medicine, CroRIS ID: 3413) based on GFP fluorescence (GFP+ for test cells and GFP- for intact or colorless UV-treated cells). The cells were sorted directly into RLT buffer to lyse them and prevent RNA degradation. Following sorting, RLT buffer lysates were frozen at −80 °C until further processing. Total cellular RNA was isolated using Qiagen RNEasy Mini kit according to manufacturer’s instructions, and DNA was removed by the addition of DNase I (New England Biolabs), also according to manufacturer’s instruction. An aliquot of DNase I treated RNA was analyzed on Agilent Bioanalyzer 2100 (Center for Proteomics, University of Rijeka Faculty of Medicine) using Eukaryote Total RNA Nano kit. All samples had RINs above 8. Isolated RNA was stored at −80 °C until sequencing libraries were generated.

### Differential Gene Expression Analysis

RNA sequencing and the following data quality control, alignment and mapping was performed by Novogene (Novogene Co., Ltd.). Quality of raw RNA-seq reads was checked using in-house script (Novogene Co., Ltd.). Clean RNA-seq reads were aligned to the *Cricetulus griseus* (Chinese hamster) reference genome (assembly GCF_000223135.1) using the Hisat2 V2.0.5 mapping tool and quantified using the featureCounts V1.5.0-p3 algorithm.

Differential gene expression analysis was performed using the DESeq2 R package v1.47.0 (45). Raw counts were first pre-filtered to remove genes with less than 10 reads in total, normalized with DESeq2 default parameters and transformed using variance stabilizing transformation (VST). To examine sample clustering, principal component analysis (PCA) was performed on the normalized counts. To correctly measure the effect of various test conditions, samples were modelled to control for batch differences and other technical differences arising from the experimental design. For effect size shrinkage, the “ashr” method was used which implements an empirical Bayes approach for large-scale hypothesis testing and false discovery rate (FDR) estimation (ashr R package v2.2-63). For further downstream analysis, differentially expressed genes (DEGs) were defined with FDR adjusted p-value < 0.05 and fold change > 2 for upregulated genes or < 0.5 for downregulated genes.

### Functional Enrichment Analysis

R package biomaRt version 2.63.0 (46,47) with the Ensembl database was used to convert gene names to Entrez ID for downstream analysis. Given the limited annotation resources for the Chinese hamster (*Cricetulus griseus*) genome, mouse (*Mus musculus*) annotations were used as a proxy. Genome wide annotation for mouse was used via org.Mm.eg.db R package v3.18.0. Functional enrichment analyses were performed with clusterProfiler R package v4.10.0 (48). Gene Ontology (GO) over-representation tests were done separately for up- and downregulated DEGs and the results were filtered based on FDR adjusted p-value < 0.05. Redundant GO terms were removed by applying semantic similarity method implemented within the function simplify, using the similarity cut-off of 0.5 (49).

### Cell communication analysis

To assess the abundance of individual ligand-receptor pairs formed between the members of two cell subpopulations from the bulk transcriptomics profiles, a novel approach ICELLNET R package v2.2.0 was used (17,50). This approach relies on a curated database comprising 1669 ligand-receptor interactions collected from the scientific literature and other public resources. These interactions are organized into ten categories of signaling molecules - Cytokines, Chemokines, Immune Checkpoints, Notch and Wnt pathways, HLA-related signals, Innate immune factors, Growth factors, Cell adhesion molecules, and ECM interactions.

ICELLNET models communication between a designated “central” (source) cell and one or more “partner” (target) cells by analysing the expression levels of genes encoding ligands and receptors. To ensure that communication scores are not disproportionately influenced by genes with high expression, DESeq2-normalized counts are rescaled before further analysis. Communication is evaluated in two directions: “outward”, where ligand is expressed in the source and receptor in the target cell, and “inward”, with the opposite direction of signalling.

Each ligand-receptor pair is assigned a score based on the product of the rescaled expression levels in the corresponding source and target cells. If either component of the interaction is not expressed, the score is zero. These interaction scores are then combined into a summary metric that represents the overall level of communication between the two cell subpopulations.

To assess cell-cell communication between intact and UV-irradiated cells, individual communication scores were first calculated for each ligand-receptor pair. The difference (delta) in these scores between the two conditions was then computed to identify the ligand-receptor pairs and the corresponding categories that were most affected. The top 10% of the most upregulated and downregulated ligand-receptor pairs were selected for more detailed analysis.

### Statistical analysis

All experiments were done in triplicates, unless indicated otherwise. Statistical tests were performed in GraphPad Prism version 8.4.3 for Windows using two-way ANOVA and Bonfferoni’s multiple comparison post-test. Exceptionally, ordinary one-way ANOVA with Bonfferoni’s post-test was used for statistical analysis of beta-galactosidase (Fig 1D) and Caspase 3/7 activity (Fig 1F). * p < 0.05; ** p < 0.01; *** p < 0.005; **** p < 0.001; ns, not significant.

Positive controls with monocultures and co-cultures were ran within all experiments with conditioned media.

### Figure Preparation

The complete bioinformatics pipeline was performed in the free software environment for statistical computing R, version 4.3.2 (R Core Team, 2023). Heatmaps were generated with the Complex Heatmap package v2.18.0. Venn diagrams were created with the eulerr package v7.0.2. For depicting results of the GO enrichment analysis clusterProfiler v4.10.0 was used (48). All other figures were created using the ggplot2 package v4.3.3. To combine individual figures the multipanelfigure package v2.1.6 was used.

## List of Abbreviations

7-AAD: 7-aminoactinomycin
D BSA: Bovine Serum Albumin
CASP3/7: Caspase-3/7
DAPI: 4’,6-diamidino-2-phenylindole
DEGs: Di?erentially Expressed Genes
DMEM: Dulbecco’s Modi?ed Eagle Medium
DMSO: Dimethyl Sulfoxide
DNase I: Deoxyribonuclease I
ECM: Extracellular Matrix
FBS: Fetal Bovine Serum
FDR: False Discovery Rate
GFP: Green Fluorescent Protein
GJ: Gap Junctions
GO: Gene Ontology
HBSS: Hank’s Balanced Salt Solution
HDF-TERT: Human Dermal Fibroblasts immortalized with telomerase
HLA: Human Leukocyte Antigen
HCl: Hydrochloric Acid
MTT: 3-(4,5-dimethylthiazol-2-yl)-2,5-diphenyltetrazolium bromide
PBS: Phosphate-Bu?ered Saline
PCA: Principal Component Analysis
PI: Propidium Iodide
PKC: Protein Kinase C
PMA: Phorbol 12-myristate 13-acetate PS Phosphatidylserine
PVDF: Polyvinylidene Fluoride
RIN: RNA Integrity Number
RNA-seq: RNA sequencing
ROS: Reactive Oxygen Species
RT: Room Temperature
SDS: Sodium Dodecyl Sulfate
Stsp: Staurosporine (apoptosis inducer)
TBS: Tris-Bu?ered Saline
TBS-T: Tris-Bu?ered Saline with Tween 20
TNTs: Tunneling Nanotubes
UV: Ultraviolet
VST: Variance Stabilizing Transformation

## Supplementary Information

Figures S1-S4 Table S1 Table S2 Table S3 Table S4 Table S5 Table S6

## Declarations

### Ethics approval and consent to participate

Not applicable.

### Consent for publication

All authors read and approved the final manuscript.

### Availability of data and materials

RNA sequencing datasets are available in the BioStudies database (http://www.ebi.ac.uk/biostudies) under accession number E-MTAB-15733. All other datasets supporting the conclusions of this article are included within the article and its additional files.

## Authors’ contributions

Conceptualization – KT, MR; Methodology – KT, SS, MN, JB; Software – AG; Validation – JB; Formal analysis – JB, NP, AG; Investigation – JB, NP, VJL, KT; Resources – KT, MR; Data curation – JB, AG; Writing – Original Draft – KT, JB; Writing – Review & Editing – AG, VJL, SS, MN, MR; Visualization – JB, AG, NP; Supervision – KT; Project administration – KT; Funding acquisition – KT, MR.

## Acknowledgements

We thank Sara Trifunovic for performing preliminary experiments, Danea Jonjic for technical assistance with quantification of the nuclear phenotypes, and Anita Krisko for critical reading of the manuscript and helpful comments.

