## Supplementary material for "Contact-dependent regulation of UV-B/C-induced cell fate by neighbouring intact cells": Figures S1-S4

### Slide 1
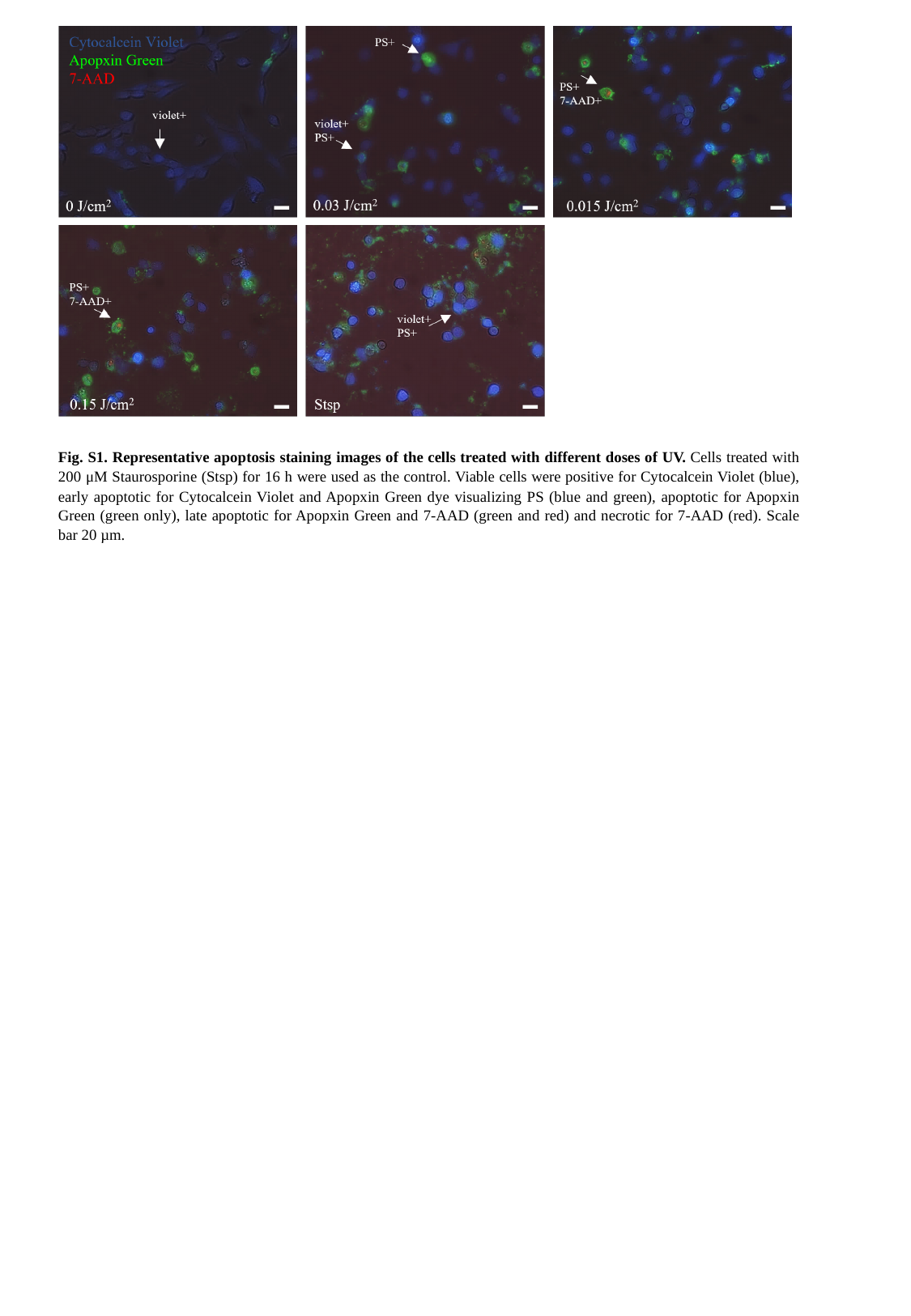

Fig. S1. Representative apoptosis staining images of the cells treated with different doses of UV. Cells treated with 200 μM Staurosporine (Stsp) for 16 h were used as the control. Viable cells were positive for Cytocalcein Violet (blue), early apoptotic for Cytocalcein Violet and Apopxin Green dye visualizing PS (blue and green), apoptotic for Apopxin Green (green only), late apoptotic for Apopxin Green and 7-AAD (green and red) and necrotic for 7-AAD (red). Scale bar 20 µm.

### Slide 2
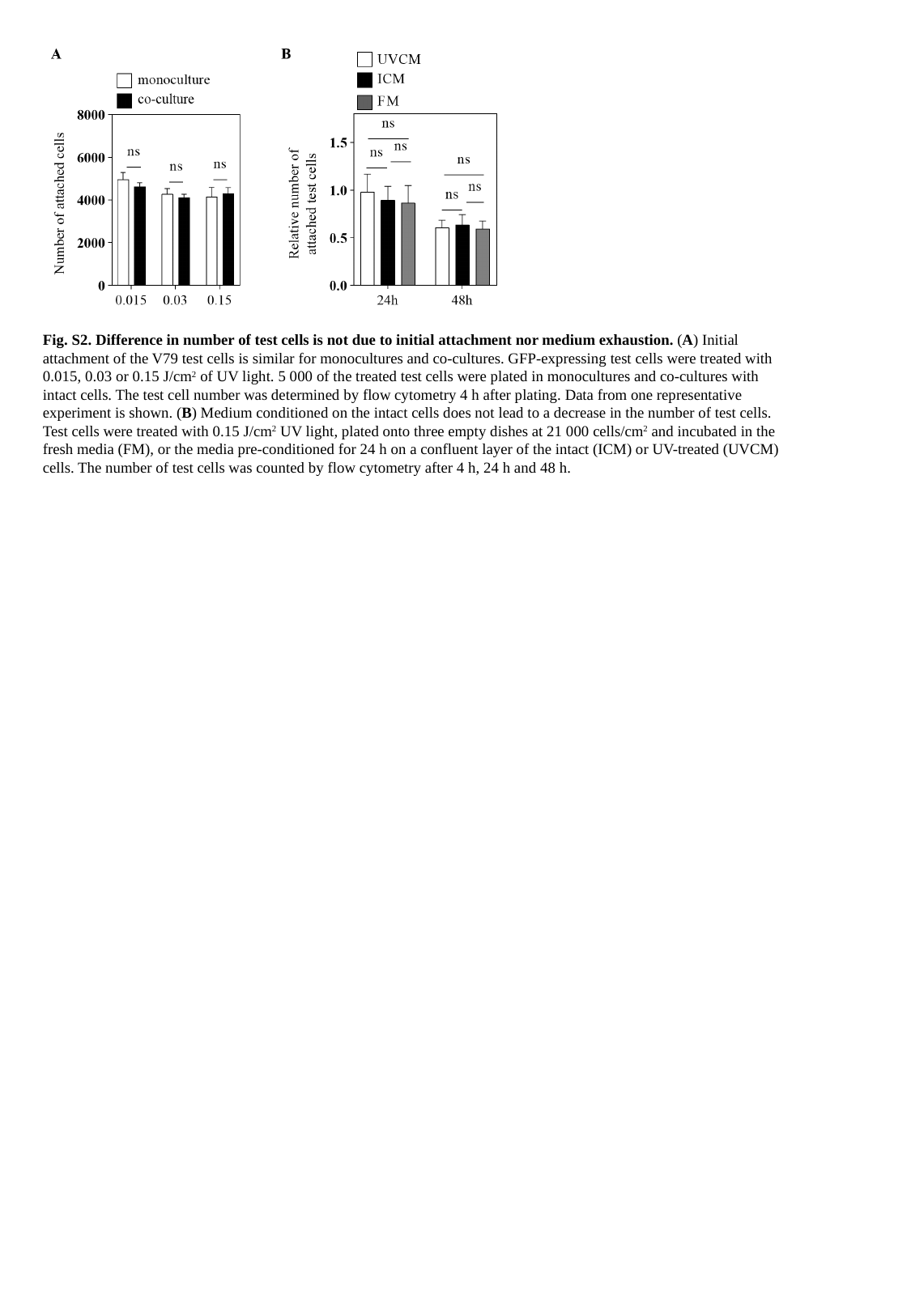

Fig. S2. Difference in number of test cells is not due to initial attachment nor medium exhaustion. (A) Initial attachment of the V79 test cells is similar for monocultures and co-cultures. GFP-expressing test cells were treated with 0.015, 0.03 or 0.15 J/cm^2^ of UV light. 5 000 of the treated test cells were plated in monocultures and co-cultures with intact cells. The test cell number was determined by flow cytometry 4 h after plating. Data from one representative experiment is shown. (B) Medium conditioned on the intact cells does not lead to a decrease in the number of test cells. Test cells were treated with 0.15 J/cm^2^ UV light, plated onto three empty dishes at 21 000 cells/cm^2^ and incubated in the fresh media (FM), or the media pre-conditioned for 24 h on a confluent layer of the intact (ICM) or UV-treated (UVCM) cells. The number of test cells was counted by flow cytometry after 4 h, 24 h and 48 h.

### Slide 3
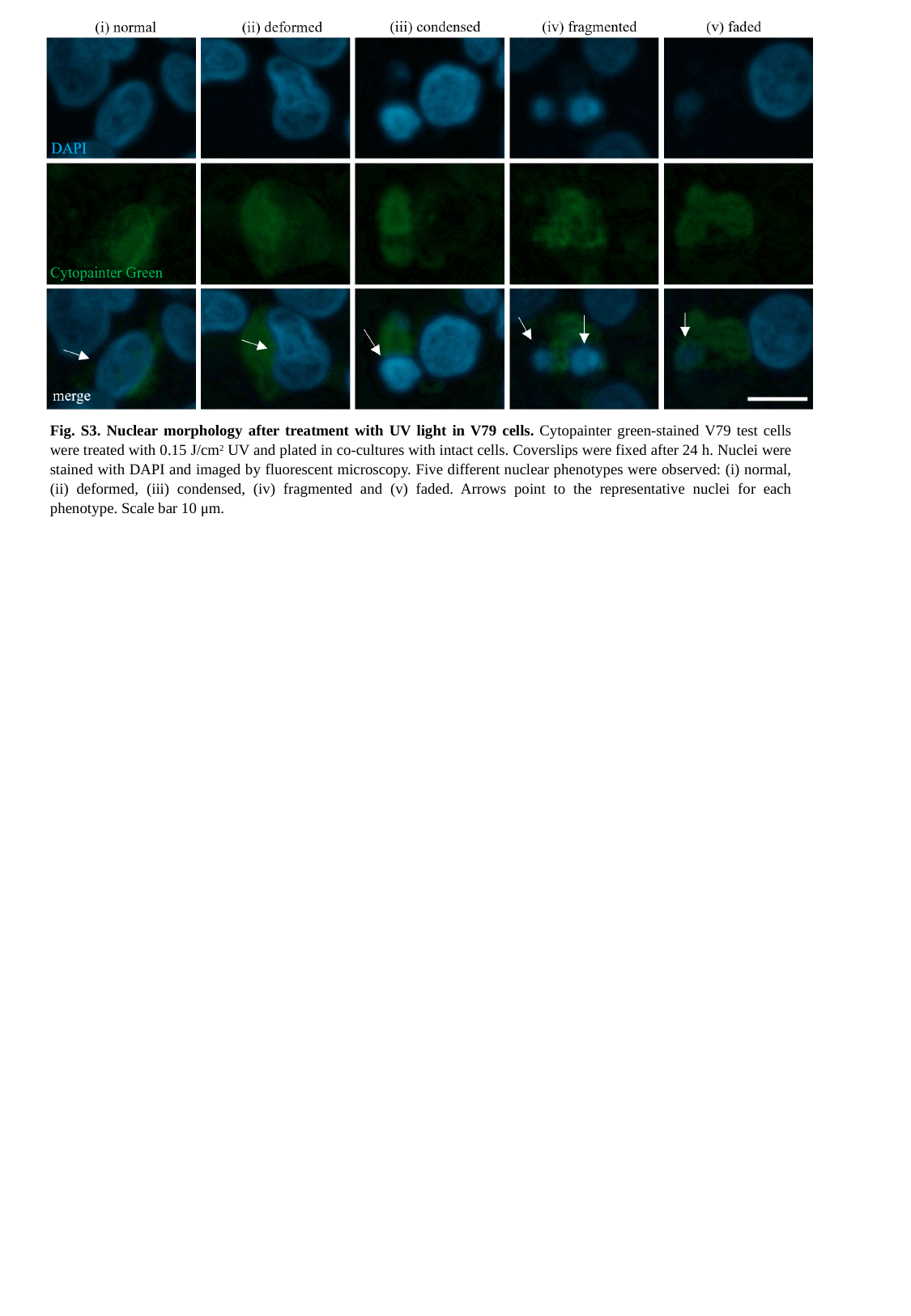

Fig. S3. Nuclear morphology after treatment with UV light in V79 cells. Cytopainter green-stained V79 test cells were treated with 0.15 J/cm^2^ UV and plated in co-cultures with intact cells. Coverslips were fixed after 24 h. Nuclei were stained with DAPI and imaged by fluorescent microscopy. Five different nuclear phenotypes were observed: (i) normal, (ii) deformed, (iii) condensed, (iv) fragmented and (v) faded. Arrows point to the representative nuclei for each phenotype. Scale bar 10 μm.

### Slide 4
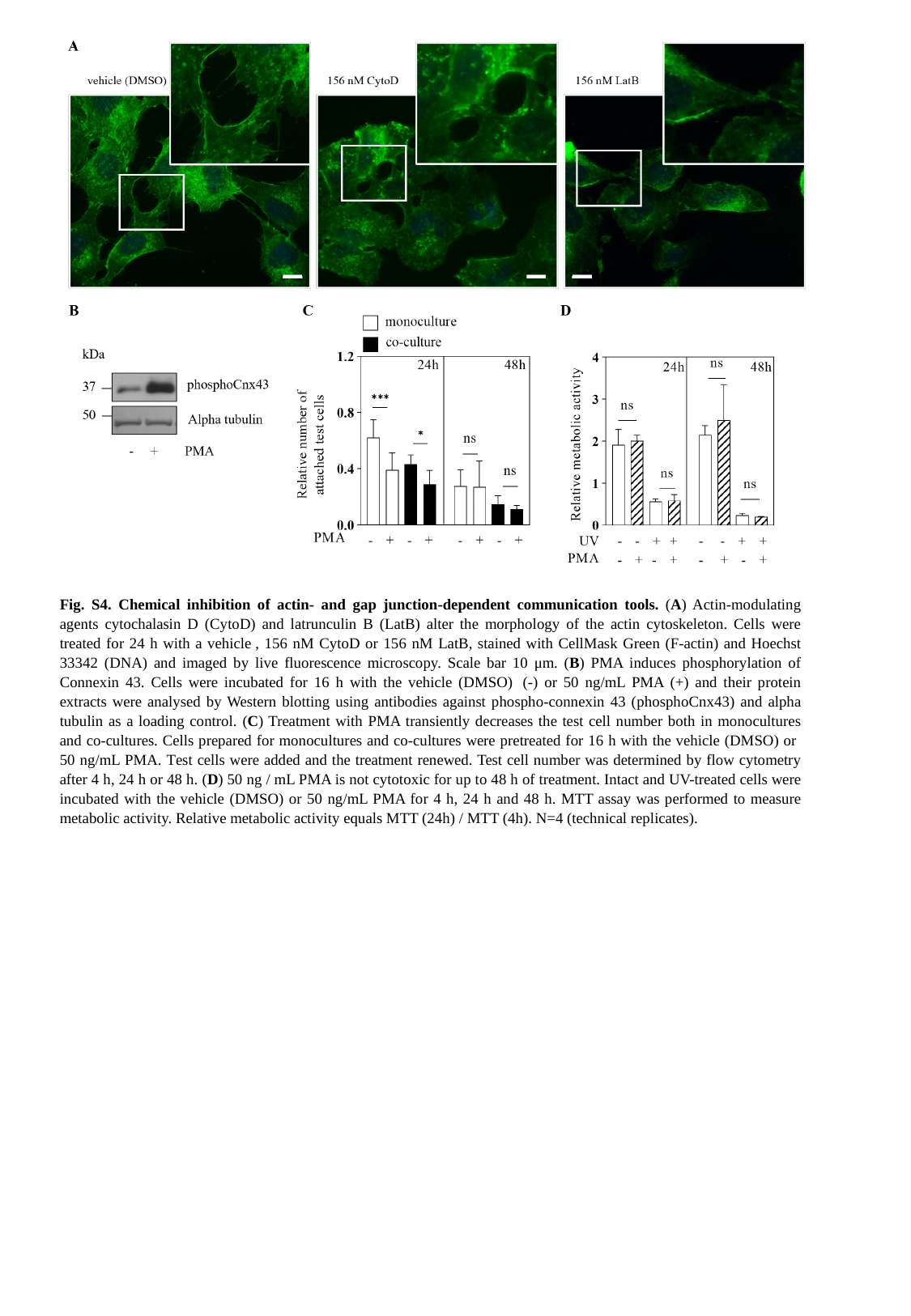

Fig. S4. Chemical inhibition of actin- and gap junction-dependent communication tools. (A) Actin-modulating agents cytochalasin D (CytoD) and latrunculin B (LatB) alter the morphology of the actin cytoskeleton. Cells were treated for 24 h with a vehicle , 156 nM CytoD or 156 nM LatB, stained with CellMask Green (F-actin) and Hoechst 33342 (DNA) and imaged by live fluorescence microscopy. Scale bar 10 μm. (B) PMA induces phosphorylation of Connexin 43. Cells were incubated for 16 h with the vehicle (DMSO)  (-) or 50 ng/mL PMA (+) and their protein extracts were analysed by Western blotting using antibodies against phospho-connexin 43 (phosphoCnx43) and alpha tubulin as a loading control. (C) Treatment with PMA transiently decreases the test cell number both in monocultures and co-cultures. Cells prepared for monocultures and co-cultures were pretreated for 16 h with the vehicle (DMSO) or 50 ng/mL PMA. Test cells were added and the treatment renewed. Test cell number was determined by flow cytometry after 4 h, 24 h or 48 h. (D) 50 ng / mL PMA is not cytotoxic for up to 48 h of treatment. Intact and UV-treated cells were incubated with the vehicle (DMSO) or 50 ng/mL PMA for 4 h, 24 h and 48 h. MTT assay was performed to measure metabolic activity. Relative metabolic activity equals MTT (24h) / MTT (4h). N=4 (technical replicates).
